# Sparcl1/Hevin drives inflammatory and neuropathic pain through astrocyte and NMDA receptor signaling

**DOI:** 10.1101/2022.03.28.486142

**Authors:** Gang Chen, Hao Luo, Xin Luo, Sandeep K. Singh, Juan J. Ramirez, Michael L. James, Joseph P. Mathew, Miles Berger, Cagla Eroglu, Ru-Rong Ji

**Affiliations:** Center for Translational Pain Medicine, Department of Anesthesiology, Duke University Medical Center, Durham, NC 27710, USA; Department of Cell Biology, Duke University Medical Center, Durham, NC 27710, USA; Department of Biochemistry and Molecular Biology, Virginia Commonwealth University School of Medicine, Richmond, VA 23298, USA; Department of Neurobiology, Duke University Medical Center, Durham, NC 27710, USA; Department of Anesthesiology, Duke University Medical Center, Durham, NC 27710, USA; Howard Hughes Medical Institute, Duke University Medical Center, Durham, NC 27710, USA; Duke Institute for Brain Sciences (DIBS), Durham, NC 27710, USA

## Abstract

Hevin/Sparcl1 is an astrocyte-secreted protein and regulates synapse formation in the brain. Here we show that astrocytic hevin signaling plays a critical role in maintaining chronic pain. Compared to wild-type mice, hevin-null mice exhibited normal mechanical and heat sensitivity but reduced inflammatory pain. Interestingly, hevin is required for the maintenance of nerve injury-induced neuropathic pain (mechanical allodynia), and hevin-null mice have faster recovery than wild-type mice from neuropathic pain after nerve injury. Intrathecal injection of wild-type hevin but not a hevin mutant that is no longer synaptogenic was sufficient to induce persistent mechanical allodynia in naïve mice and further enhanced neuropathic pain in animals with nerve injury. In hevin-null mice with nerve injury, AAV-mediated re-expression of hevin, but not mutant hevin, in GFAP-expressing spinal cord astrocytes could reinstate neuropathic pain. Mechanistically, hevin is crucial for spinal cord NMDA receptor (NMDAR) signaling, as NMDA-induced mechanical allodynia and inward currents in spinal cord lamina II neurons is reduced in hevin-null mice. Hevin potentiated NMDA currents mediated by the GluN2B-containing NMDARs. Furthermore, intrathecal injection of a neutralizing antibody against hevin alleviated acute inflammatory pain and persistent neuropathic pain. Secreted hevin was detected in mouse cerebrospinal fluid (CSF) and nerve injury significantly increased CSF hevin abundance. Finally, neurosurgery caused rapid (< 10 hours) and substantial increases (~20 fold) in HEVIN levels in human CSF. Collectively, our findings support a critical role of hevin and astrocytes in the maintenance of chronic pain. Neutralizing of secreted hevin with monoclonal antibody may provide a new therapeutic strategy for treating chronic pain and NMDAR-medicated neurodegeneration.

---

Chronic pain, such as neuropathic pain is a major health problem worldwide and difficult to treat (1, 2). It is generally believed that maladaptive synaptic plasticity in the spinal cord and brain drives chronic pain (1, 3). Activation of NMDA receptors, especially NR2B subunit (GluN2B) plays a crucial role in injury-induced synaptic plasticity and pain pathogenesis (4, 5). Astrocytes are a major glial cell type in the central nervous system (CNS) and critical for maintaining CNS homeostasis, including physiological pain (6–8). Astrocytes make close contacts with synapses, and therefore, are positioned to regulate synapse formation and synaptic plasticity (9–12). During development astrocytes promote synapse formation and regulate synaptic connectivity in the CNS through secreted signals, such as thrombospondin-4 (TSP4) (9, 10, 13) or adhesion molecules, such as neuroligins and hepatocyte cell adhesion molecule (11, 14). Multiple types of neurological and neuropsychiatric disorders, such as Alzheimer’s disease and chronic pain, may result from ‘gliopathy’ of astrocytes (15), leading to an inflammatory and pathological A1-like phenotype (16). Several lines of evidence support an essential role of astrocytes in neuropathic pain development and maintenance (15, 17–19). Following painful nerve injury or chemotherapy-induced neuropathy, astrogliosis is more prominent and persistent than microgliosis in the spinal cord dorsal horn (SDH) (15, 20). Several signal molecules, such as connexin-43 (Cx43), and chemokines (CCL2 and CXCL1), MAP kinases (ERK/JNK), and JAK-STAT3 have been implicated in astrocyte signaling to maintain neuropathic pain (6, 18, 19, 21). Nerve injury upregulates TSP4 in spinal astrocytes and facilitates neuropathic pain via modulation of excitatory synaptic transmission (22). Gabapentin is a widely used drug for treating neuropathic pain through interaction with calcium channel alpha2delta-1 subunit (Cavα2δ1), but interestingly, Cavα2δ1 has been identified as a thrombospondin receptor and regulates excitatory synaptogenesis (13). TSP4-induced pain can be blocked by gabapentin or Cavα2δ1 knockdown (23). Furthermore, a population of astrocytes in superficial SDH is involved in descending noradrenergic control of mechanical pain (24). However, it is not fully understood how astrocytes regulate synaptic plasticity in pathological pain.

<u>Hevin,</u> short for high endothelial venule protein, also known as SPARC-like 1 (SPARCL1) or synaptic cleft 1, SC1 (25), is a member of the SPARC family of glycoproteins that regulate cell-matrix interactions. Like TSP4, hevin is an astrocyte-secreted synaptogenic protein (9, 10), but its role in chronic pain is unclear. Using a combination of in vitro and in vivo approaches, we demonstrated that hevin assembles Vesicular Glutamate Transporter 2 (VGlut2) positive thalamocortical synapses by bridging neurexin-1alpha (Nrx1α, a presynaptic component) and neuroligin-1B (NL1B, a postsynaptic component) (9). Neurexin-1alpha and NL1B do not interact with each other under normal conditions; but hevin can bridge these two adhesion proteins across the synapse, through interactions mapped to a specific synaptogenic domain (9). These interactions of hevin are critical for the formation and plasticity of thalamocortical synapses in the developing visual cortex via specific regulation of GluN2B-containing NMDARs (9). In this study, we investigated the role of hevin in physiological, inflammatory, and neuropathic pain. We demonstrated that hevin is sufficient and required for inducing central sensitization (synaptic plasticity in the spinal cord pain circuit) and mechanical pain (tactile allodynia) via NMDAR/GluN2B signaling. Our data also show that hevin is secreted in mouse and human CSF under injury conditions, and neutralization of the secreted hevin protein with a monoclonal antibody effectively alleviates neuropathic pain. Furthermore, we demonstrate that re-expression of hevin in spinal cord astrocytes of knockout mice lacking hevin is sufficient to reinstate neuropathic pain.

## Results

### Hevin Is Distinctly Required in Physiological Pain, Inflammatory Pain, and Neuropathic Pain

We first set out to assess pain sensitivity in wild-type (WT) and hevin-knockout (KO) mice in physiological and pathological conditions. Compared to WT control, KO mice exhibited normal baseline mechanical sensitivity as measured in von Frey testing (**Fig. 1*A***) and thermal sensitivity in Hargreaves test, hot plate, and tail immersion tests (**Fig. 1 *B-D***), suggesting that hevin is dispensable for physiological pain. Next, we examined formalin-induced acute inflammatory pain, which is typically divided into phase-1 (0-10 min) and phase-2 (10-45 min). Notably, phase-2 spontaneous pain is driven by NMDAR-mediated spinal neuron sensitization (central sensitization) (26). We found that formalin-induced phase-2, but not phase-1 spontaneous pain was significantly reduced in KO mice (**Fig. 1*E***), supporting a role of hevin in central sensitization-mediated pain. Intraplantar carrageenan induced rapid and persistent inflammatory pain that fully recovered in 3 days in WT mice; but hevin-KO mice showed a faster recovery of this inflammatory pain (**Fig. 1*F***). Chronic constriction injury (CCI) of the sciatic nerve induced neuropathic pain, manifested as mechanical allodynia, a reduction in paw withdrawal threshold (PWT), which was maintained at 28 d in WT mice. Strikingly, even though the induction phase of neuropathic pain (5-10 d) was not altered in KO mice compared to WT mice, the CCI-induced mechanical allodynia was fully recovered at 28 d in hevin-KO mice, (**Fig. 1*G***). We also measured CCI-induced ongoing pain (3 w after CCI) using a 2-chamber conditioned place preference test (27). The result showed that hevin-KO mice spent less time than WT mice in the analgesic clonidine-treated chamber (**Fig. 1*H***), indicating that nerve injury-induced ongoing pain maintenance is reduced in hevin-KO mice. These results show that hevin is required for driving inflammatory and neuropathic pain but dispensable for physiological pain.

**Fig. 1.**
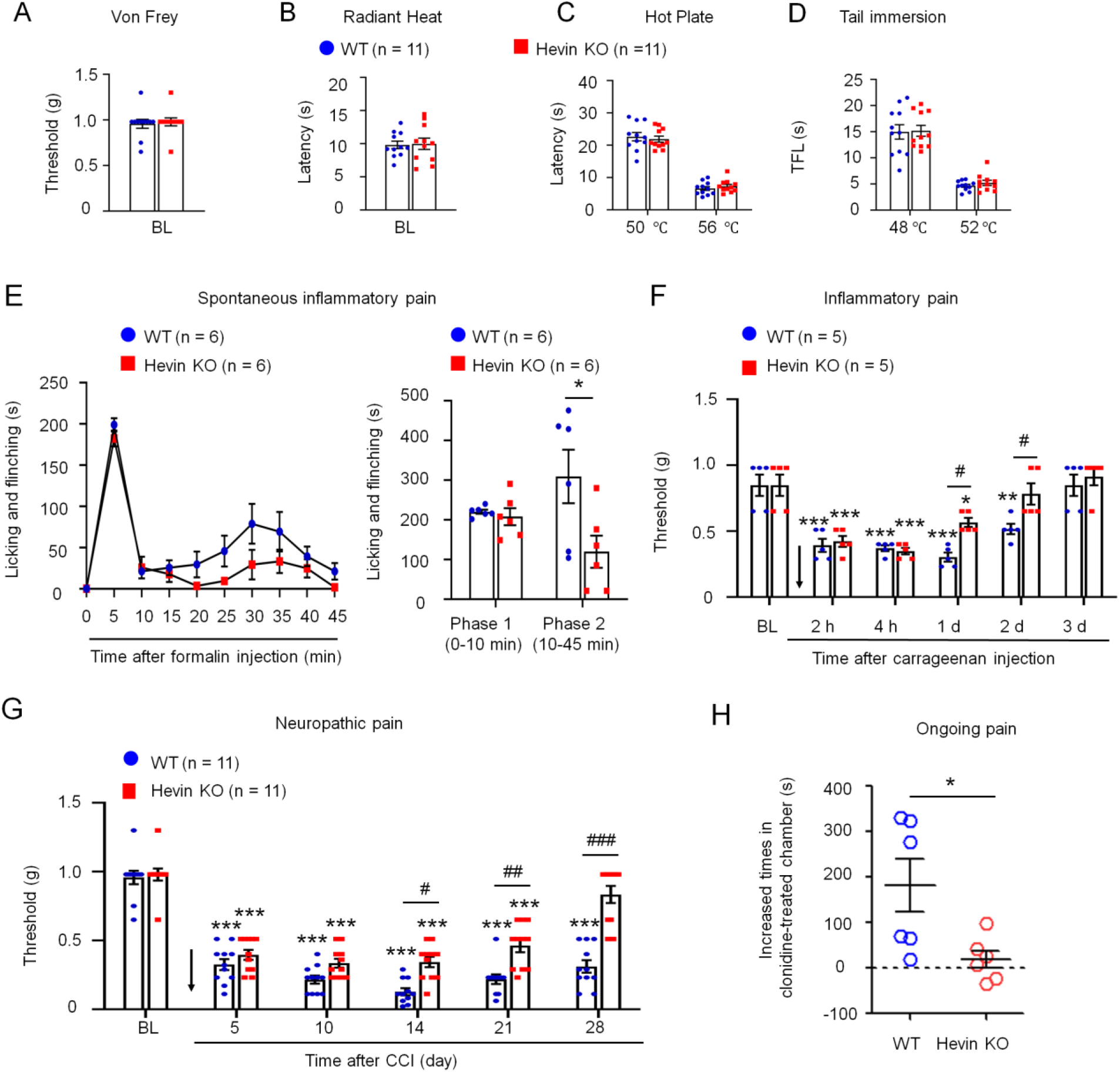
Baseline pain, inflammatory pain, and neuropathic pain in WT and hevin-KO mice. (A-D) There are no significant differences in mechanical and thermal pain threshold between WT and hevin-KO male mice, as shown in von Frey test (A), radiant heat test (B), hot plate test (C) and tail immersion test (D). *P* > 0.05, unpaired t-test, n = 11 mice/group. (E) Formalin-induced acute inflammatory pain was significantly reduced in hevin-KO male mice. Left, time-course of licking and flinching behavior following intraplantar injection of 5% formalin. Right, formalin-induced Phase I (1–10 min) and Phase II (10–45 min) responses. \**P* < 0.05, Two-way ANOVA followed by Bonferroni posthoc test. n = 6 mice/group. (F) Mechanical allodynia, induced by intraplantar injection of carrageenan, recovered faster in hevin-KO male mice than in WT male mice. Arrow indicates the time of carrageenan injection. \**P* < 0.05, \*\**P* < 0.01, \*\*\**P* < 0.001, compared to BL group, #*P* < 0.05, Two-way ANOVA followed by Bonferroni posthoc test. n = 5 mice/group. Data shown as mean ± SEM. (G) Hevin-KO male mice recovered faster from CCI-induced persistent mechanical allodynia than WT male mice. \*\*\**P* < 0.001, compared to BL group; #*P* < 0.05, ##*P* < 0.01, ###*P* < 0.001, Two-way ANOVA followed by Bonferroni posthoc test. n = 11 mice/group. (H) Ongoing pain 3 weeks after CCI in WT and hevin-KO male mice were tested using a 2-chamber conditional place preference. Ongoing pain was present in WT mice but absent in hevin-KO mice following clonidine treatment (10 μg, i.t.). \**P* < 0.05, unpaired t-test, n = 6 mice/group. All data are shown as mean ± SEM.

### Hevin Is Sufficient to Induce Mechanical Allodynia in WT Mice

Next, we evaluated whether administration of exogenous hevin, via intrathecal route to target spinal cord, in naïve mice would elicit mechanical pain. We also included a hevin mutant which cannot bridge Nrx1a and NL1B thus cannot induce synapse formation (hevin-ΔDE, lacking a.a. 351-440, *SI Appendix*, Fig. S1) as a negative control (9). Intrathecal injection of purified WT hevin (10 μg, i.t.) but not hevin-ΔDE (10 μg, i.t.) induced robust and persistent mechanical allodynia in naïve male mice, this effect lasted for more than 3 days and recovered after 5 days (**Fig. 2*A***). We also examined whether hevin could exacerbate neuropathic pain. Intrathecal injections of WT but not hevin-ΔDE, given 12 d after CCI, further enhanced mechanical allodynia in male mice with nerve injury (**Fig. 2*B***). These results suggested that hevin is sufficient to induce mechanical allodynia in naïve mice and further enhance pain in neuropathic pain mice. We did not find sex differences in hevin-induced pain, as i.t. hevin induced robust mechanical allodynia in both males and females (*SI Appendix*, Fig. S2a).

**Fig. 2.**
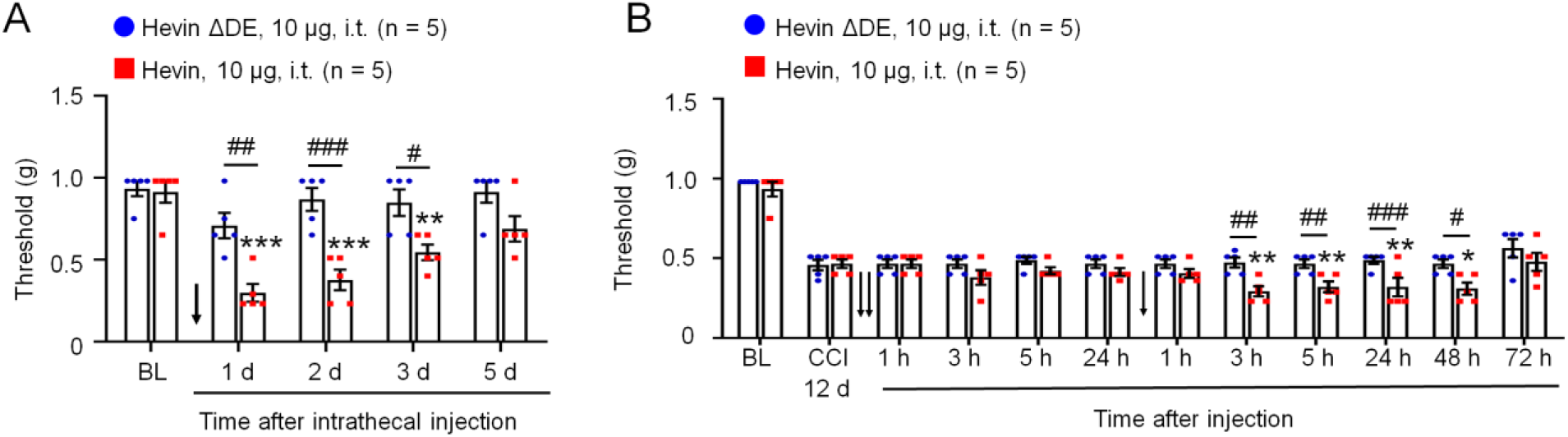
Intrathecal administration of hevin but not hevin-ΔDE (10 μg, i.t.) decreases paw withdrawal threshold in naïve mice and mice with nerve injury. (A) Intrathecal injection of WT hevin but not mutant hevin (hevin-ΔDE) induced mechanical allodynia in naïve mice, lasting > 3 days. n = 5 mice/group. \*\**P* < 0.01, \*\*\**P* < 0.001, compared to BL group, #*P* < 0.05, ##*P* < 0.01, ###*P* < 0.001, Two-way ANOVA followed by Bonferroni posthoc test. Arrow indicates hevin injection on day 0. (B) Repeated intrathecal injections of WT hevin but not mutant hevin (hevin-ΔDE), given at 12 and 13 days after CCI induced, further exacerbated mechanical allodynia. n = 5 mice/group. \**P* < 0.05, \*\**P* < 0.01, compared to CCI-12d baseline group; #*P* < 0.05, ##*P* < 0.01, ###*P* < 0.001, Two-way ANOVA followed by Bonferroni posthoc test. Arrows indicate the time of hevin injections. All data are shown as mean ± SEM.

### Re-expression of hevin In Spinal Astrocytes Reinstates Neuropathic Pain

To investigate the cellular mechanisms by which hevin modulates pain, we examined hevin expression in the mouse spinal cord dorsal horn (SDH) and found that hevin is expressed by the majority of astrocytes (GFAP^+^, **Fig. 3*A***) and some neurons (NeuN^+^, fig. S3a), but not by microglia (Iba1^+^, *SI Appendix*, Fig. S3b) in SDH. The specificity of the hevin antibody was validated by loss of hevin staining in SDH of hevin-KO mice (**Fig. 3*B***). Western blot result also confirmed hevin expression in WT-SDH, which was lost in SDH of KO mice (SI Appendix, Fig. S3c).

**Fig. 3.**
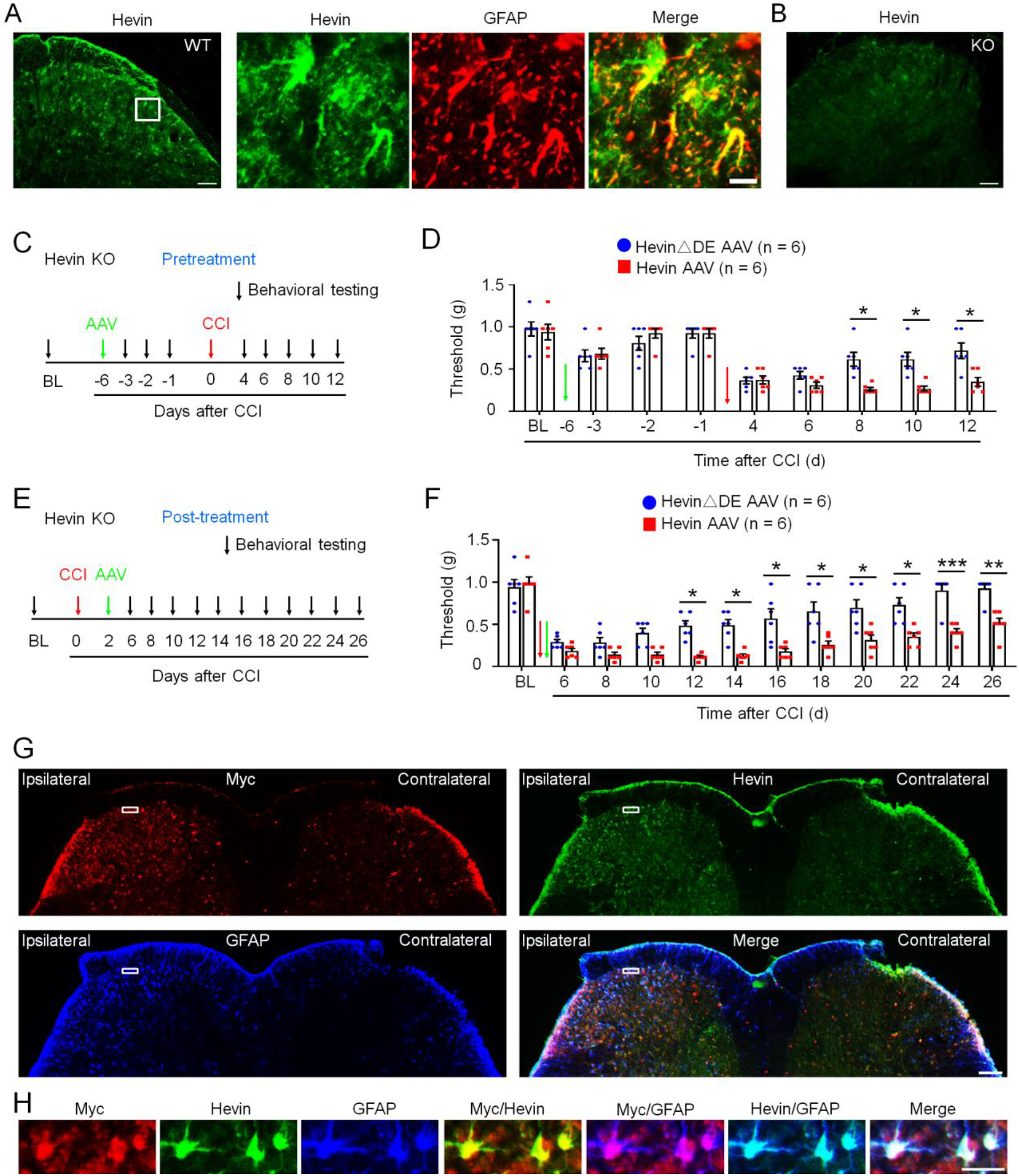
Expression of hevin in spinal astrocytes by intraspinal hevin-AAV injection can reinstate neuropathic pain in hevin KO mice. (A) Double immunostaining of hevin (green) and the astrocyte marker GFAP (red) in the spinal cord dorsal horn in mice. Note hevin is primarily colocalized with GFAP. Scale bar: 100 μm (left) and 20 μm (right). The box in the left is enlarged in the right panels. (B) Absence of hevin immunostaining in SDH in hevin KO mice. Scale, 100 μm. (C) Paradigm for measuring mechanical allodynia in hevin KO mice with intra-spinal microinjection of hevin-AAV and hevinΔDE-AAV, given 6d before CCI. (D) SDH microinjection of AAV induced mild and transient reduction in paw withdrawal threshold in naïve hevin KO mice, and there was no statistical difference between hevin-AAV and hevinΔDE-AAV groups. After CCI, mechanical allodynia was significantly prolonged in hevin-AAV treated mice than hevinΔDE-AAV treated mice. n = 6 mice/group. \**P* < 0.05, Two-way ANOVA followed by Bonferroni posthoc test. Green and red arrows indicate the time of virus injection and nerve injury, respectively. (E) Paradigm for measuring mechanical allodynia in hevin KO mice with intra-spinal microinjection of hevin-AAV and hevin-ΔDE-AAV, given 2 days after CCI. (F) SDH microinjection of hevin-AAV, given after nerve injury, significantly enhanced and prolonged mechanical allodynia in hevin KO mice compared to the hevinΔDE-AAV treated mice. n = 6 mice/group. \**P* < 0.05, \*\**P* < 0.01, \*\*\**P* < 0.001, Two-way ANOVA followed by Bonferroni posthoc test. Arrows indicate the time of virus injection and nerve injury. (G) Triple immunostaining of Myc (red, note that hevin-AAV contains a Myc-tag), hevin (green) and GFAP (blue) in the spinal cord dorsal horn in hevin KO mice, 24 days after the ipsilateral SDH hevin-AAV injection. Note hevin expression is absent in the contralateral SDH of hevin-KO mice. Scale, 100 μm. (H) Enlarged images in the box of F panel G, with additional merged images for Myc/hevin, Myc/GFAP, and hevin/GFAP. Note that hevin-expressing astrocytes also express Myc and GFAP in superficial SDH. Scale, 20 μm. All data are shown as mean ± SEM.

To determine a specific role of hevin from spinal astrocytes in neuropathic pain, we evaluated whether hevin re-expression in SDH astrocytes would reinstate neuropathic pain in hevin-KO mice (**Fig. 2G**). To this end, we conducted SDH microinjection of adeno-associated viruses (AAV2/5.GFAP.Hevin-MycHis, termed as hevin-AAV, SI Appendix, Fig. S1b) (9) to re-express hevin in spinal astrocytes in hevin-KO mice. We also included AAV2/5.GFAP.HevinΔDE-MycHis virus as a negative control (termed as hevinΔDE-AAV, fig. S1b). AAVs were injected to the ipsilateral SDH of KO mice either prior to CCI nerve injury (pretreatment, **Fig. 3 *C* and *D***) or after CCI (post-treatment, **Fig. 3 *E* and *F***). The pre-treatment in naïve mice induced mild and transient reduction in PWT in both hevin-AAV and hevinΔDE-AAV groups, due to spine surgery; but this PWT change fully recovered 5 days after the surgery (**Fig. 3*D***). Following CCI, mice treated with hevinΔDE-AAV exhibited similar time course of PWT change as in hevin-KO mice (**Fig. 2*G***), and neuropathic pain began to recover on CCI-8d (**Fig. 3*D***). Strikingly, hevin-AAV-treated KO mice showed no sign of neuropathic pain recovery on CCI 8-12d, and mechanical allodynia was significantly prolonged compared to hevinΔDE-AAV treated mice (*P* < 0.05, **Fig. 3*D***). Furthermore, post-treatment of hevin-AAV, given 2 days after CCI (**Fig. 3*E***), also significantly decreased PWT compared to the hevinΔDE-AAV mice (*P* < 0.05), displaying prolonged neuropathic pain on CCI-26d (**Fig. 3*F***). Thus, both pre- and post-treatments of hevin-AAV in KO mice restore neuropathic pain by enhancing and sustaining mechanical allodynia after CCI.

We confirmed the specific expression of hevin in SDH astrocytes after the hevin-AAV injection, in the spinal cord sections of hevin-KO mice. We used antibodies against hevin and GFAP, as well as Myc (**Fig. 3, *G* and *H***), as hevin-AAV has a Myc-tag (*SI Appendix*, Fig. S1b). At 24 d after the ipsilateral SDH hevin-AAV injection, we found many GFAP+ astrocytes in the ipsilateral SDH which were also labeled for hevin and Myc (**Fig. 3*G***), especially laminae I-III, a critical region for the transmission of mechanical pain (28, 29). High magnification images further revealed that hevin+ astrocytes also expressed Myc and GFAP in superficial SDH (**Fig. 3*H***). We did not observe hevin expression in the contralateral SDH of hevin-KO mice (**Fig. 3*G***). As previously reported (30), CCI caused a dramatic increase in the number of GFAP+ cells (astrogliosis) in the ipsilateral SDH. Collectively, these behavioral and histochemical data suggest that re-expression of hevin in spinal cord astrocytes is sufficient to reinstate neuropathic pain.

### Hevin Regulates NMDA-Evoked Pain and NMDA Currents in SDH Neurons

Previously we found that hevin strongly enhances NR2B subunit-containing NMDA receptor function by increasing NMDA currents in autoptic neuron-only cultures (9). Thus, here we tested the function of spinal NMDARs in WT and hevin-KO mice using behavioral and electrophysiological approaches. Spinal injection of NMDA (3 nmol, i.t.) induced persistent mechanical allodynia in WT mice, which lasted for >2 weeks and recovered after 3 weeks (**Fig. 4*A***). The duration of NMDA-evoked mechanical allodynia was remarkably shortened in hevin-KO mice, showing a full recovery at 6 days post injection (**Fig. 4*A***).

**Fig. 4.**
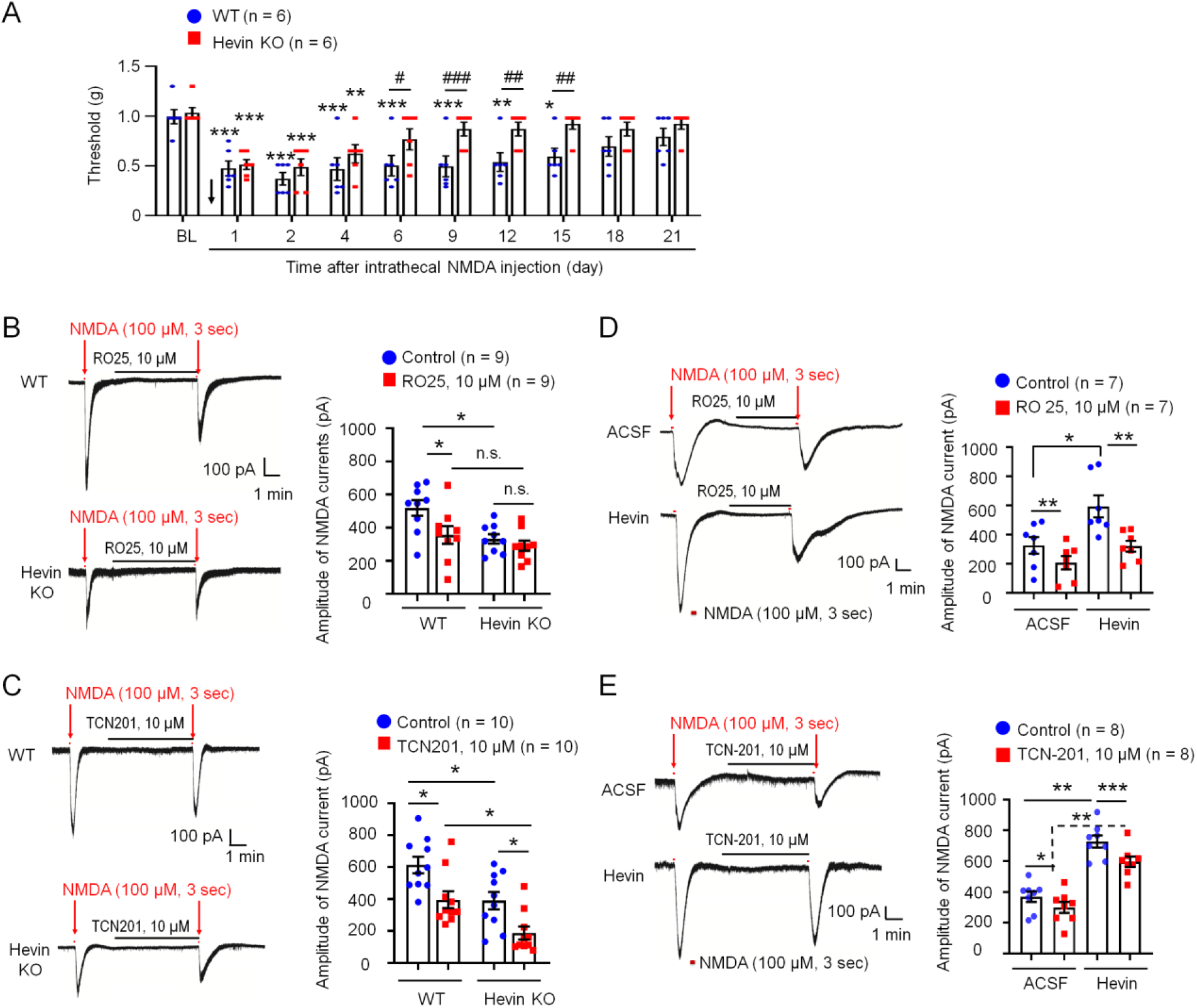
Hevin regulates NMDA-evoked pain and enhances NMDA currents in SDH neurons. (A) The duration of mechanical allodynia, induced by intrathecal injection of NMDA (3 nmol), is significantly shorter in hevin KO mice than in wild-type mice. n = 6 mice/group. \**P* < 0.05, \*\**P* < 0.01, \*\*\**P* < 0.001, compared to BL group; #*P* < 0.05, ##*P* < 0.01, ###*P* < 0.001, Two-way ANOVA followed by Bonferroni posthoc test. Arrow indicates the time of NMDA injection. (B, C) Left: representative traces of inward currents in WT and hevin KO mice, induced by NMDA (100 μM, 3 sec) via bath application. Note smaller NMDA currents in hevin KO mice and different effects of Ro25-6091 (GluN2B antagonist) and TCN-201 (GluN2A antagonist). Right: amplitude of NMDA-induced currents. n = 9, 10 neurons per group (shown in each column). \**P*<0.05, One-Way ANOVA followed by Bonferroni posthoc test. n.s., not significant. (D, E) Left: representative traces of inward currents in ACSF and hevin (100 ng/ml, 4 min) treated spinal cord slices, induced by NMDA (100 μM, 3 sec) via bath application. Note different effects of Ro25-6091 and TCN-201. Right: amplitude of NMDA-induced currents. n = 7 neurons per group. \**P* < 0.05, \*\**P* < 0.01, \*\*\**P* < 0.001, One-Way ANOVA followed by Bonferroni posthoc test. n.s., not significant. Data are shown as mean ± SEM.

We also compared NMDA-induced currents in spinal lamina IIo neurons via bath application of NMDA (100 μM, 3 sec) to spinal cord slices in WT and hevin-KO mice. Compared to WT neurons, hevin-KO neurons had significantly smaller NMDA-elicited currents (*P* < 0.05, **Fig. 4*B***). Since GluN2A and GluN2B containing NMDARs may play different roles in synaptic transmission and synaptic plasticity (5, 26), we further tested the blocking effects of GluN2A antagonist TCN201 (10 μM) and GluN2B antagonist RO25 (10 μM) on NMDA currents in spinal cord slices from WT and hevin-KO mice. RO25 produced a significant inhibition of NMDA currents in WT neurons (31%, *P*<0.05) but only evoked mild inhibition (12%, p>0.05) of NMDA currents in hevin-deficient neurons (**Fig. 4*B***). In contrast, TCN (10 μM) produced significant and comparable inhibition of NMDA currents in both WT (26%, *P* < 0.05) and KO (26%, *P* < 0.05) neurons (**Fig. 4*C***). Therefore, we conclude that hevin deficiency primarily affects the GluN2B-mediated NMDA currents in lamina IIo neurons.

Next, we investigated whether perfusion of spinal cord slices with hevin is sufficient to enhance NMDA-evoked currents in WT mice. A brief exposure of SDH neurons to hevin (4 min, 100 ng/ml ≈ 0.14 nM) elicited a significant increase in NMDA-induced currents (Before: 450.3 ± 75.5 pA; After: 625.9 ± 87.3 pA, *P* < 0.05, *SI Appendix*, Fig. S4a). Hevin-induced potentiation of NMDA currents was completely blocked by GluN2B antagonist RO25 (10 μM, **Fig. 4*D***). In contrast, GluN2A antagonist TCN (10 μM) produced a similar inhibition of NMDA currents in ACSF and hevin treated neurons (19% vs. 18% inhibition, *P* < 0.05, **Fig. 4*E***). Hevin still evoked significant increase in NMDA currents after TNC-201 treatment (*P* < 0.05, **Fig. 4*E***). However, hevin had no effects on AMPA-evoked currents in lamina IIo neurons (fig. S4b). We further assessed evoked excitatory postsynaptic currents (eEPSCs) using dorsal root stimulation. Hevin enhanced the NMDA receptor–mediated eEPSCs in lamina IIo neurons (*SI Appendix*, Fig. S4c). Collectively, these data suggest that hevin is sufficient and required to regulate GluN2B/NMDAR-mediated currents in SDH neurons. This hypothesis was further supported by our behavioral observation that mechanical allodynia by intrathecal hevin was completely reversed by RO25 in both sexes (*SI Appendix*, Fig. S2b).

### Hevin–Neutralizing Antibody Reduces Inflammatory and Neuropathic Pain in WT Mice

To examine if hevin is secreted from the mouse spinal cord, we measured hevin levels in the cerebrospinal fluid (CSF) using a mouse hevin-specific ELISA kit. We detected a high basal level of hevin secretion (~5 ng/ml) in the CSF of naïve mice (**Fig. 5*A***). To determine the role of secreted hevin in inflammatory and neuropathic pain, we tested a monoclonal antibody against mouse hevin which we previously characterized (9). The monoclonal antibody recognizes an epitope mapping to the amino acids 368–419 of hevin (anti-hevin 12:54, fig. S1c) and this antibody blocks hevin’s synaptogenic function, likely by interfering hevin’s ability to bind Nrx1a and/or NL1B (9). Another hevin antibody recognizing a different epitope (anti-hevin 12:155, fig. S1c) does not impact hevin’s synaptogenic activity, which we used as our control. Intrathecal injection of anti-hevin 12:54 (10 μg) significantly reduced formalin-induced phase II pain, compared with anti-hevin 12:155 treated mice (*P*<0.05, **Fig. 5*B***). Furthermore, intrathecal injection of anti-hevin 12:155 (10 μg), but not anti-hevin 12:155 (10 μg), rapidly (< 1h) reversed mechanical allodynia for >5h in CCI mice, in both early-phase (7d after CCI, **Fig. 5*C***) and late-phase (21d after CCI, **Fig. 5*D***) of neuropathic pain. ELISA analysis showed that CCI resulted in significant increase in CSF hevin levels (Sham: 5.24 ± 0.57 ng/ml; CCI-14d: 8.32 ± 0.89 ng/ml, *P* < 0.05, **Fig. 5*A***). These results suggest that 1) hevin is secreted in physiological and pathological conditions and 2) targeting secreted hevin with a neutralizing antibody can potently alleviate inflammatory and neuropathic pain.

**Fig. 5.**
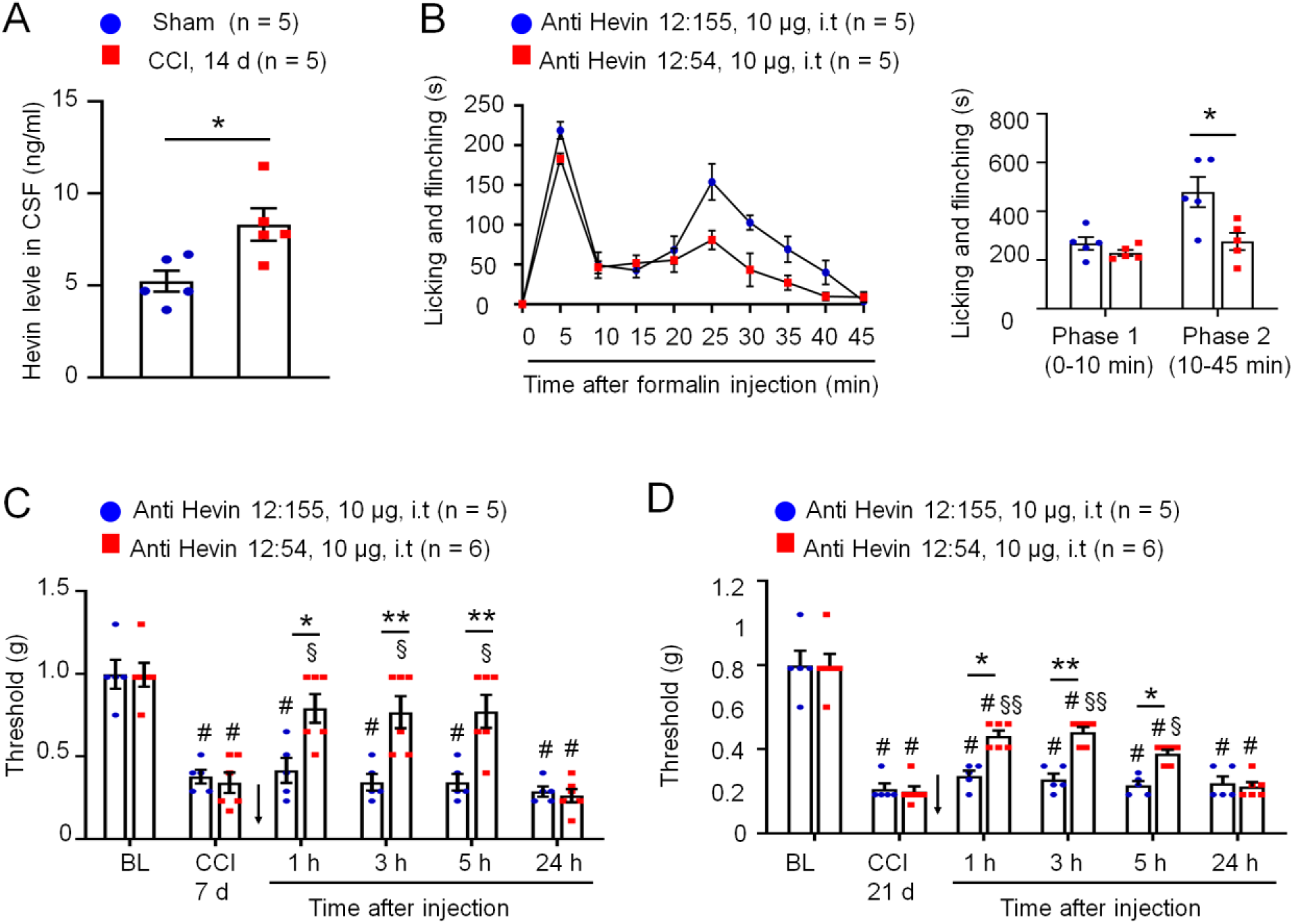
Anti-hevin monoclonal antibody 12:54 reduces inflammatory and neuropathic pain in WT mice. (A) ELISA analysis showing increased hevin level in the CSF 14 days after CCI. n = 5 mice/group. \**P* < 0.05, unpaired t-test. (B) Left, time course of formalin-induced pain in WT male mice treated with intrathecal anti-hevin 12:155 monoclonal antibody (control antibody, 10 μg) or anti-hevin 12:54 monoclonal antibody (function blocking antibody, 10 μg). n = 5 mice per group. Right, formalin-induced Phase-1 and Phase-2 responses. \**P*<0.05, unpaired t-test. (C, D) Intrathecal injection of anti-hevin 12:54 antibody (10 μg), given 7 days (C) and 21 days (D) after nerve injury, reduced CCI-induced mechanical allodynia for 5h. Arrows indicate the time of antibody injection. n = 5–6 mice/group. #*P*<0.001, versus corresponding BL group; §*P*<0.01, §§ *P*<0.001, versus corresponding baseline at CCI 7d or CCI 21d; \**P*<0.05, \*\**P*<0.01, Two-Way ANOVA followed by Bonferroni posthoc test. Data are shown as mean ± SEM.

### Hevin Is Increased in Human CSF Following Neurosurgery

To explore the translational potential of these findings, next we assessed secreted hevin levels in human CSF samples collected prior to and 10 h after neurosurgery/otolaryngology procedures (e.g., trigeminal neuralgia) (31), using a human hevin-specific ELISA kit. In the human CSF, we were able to detect a basal secretion of hevin, ranging from 0.8-2.7 ng/ml (**Fig. 6*A***). Neurosurgery resulted in a rapid (< 10 h) and dramatic increase in CSF hevin levels, ranging from 3.1-52.9 ng/ml at 10 h (**Fig. 6*A***). Strikingly, hevin showed marked increases in all the CSF samples we analyzed, ranging from 2.3-50.9 fold (*P* <0.01, **Fig. 6*B***). We also measured total protein levels in these CSF samples and observed mild but significant increases, ranging from 1.3-4.3 fold (*P*< 0.001, **Fig. 6*C***). After normalization with respective protein changes, CSF-hevin levels still exhibited a significant increase (9.0 fold, *P* <0.01, **Fig. 6*D***). These results suggest that human neurosurgery selectively increased the CSF section of hevin, beyond the postoperative increase in CSF total protein levels.

**Fig. 6.**
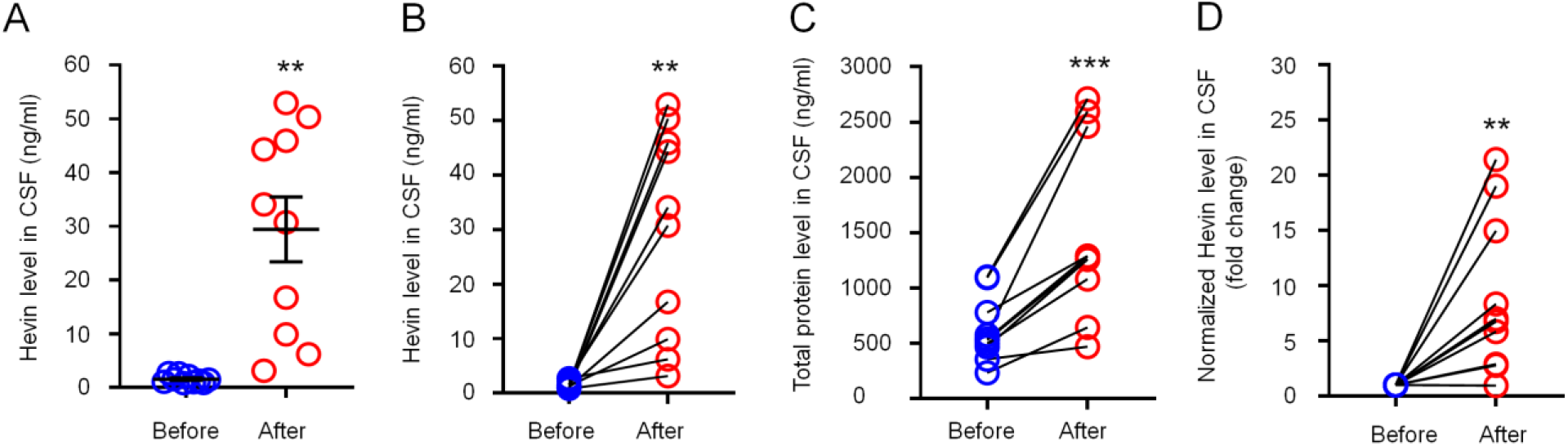
Neurosurgery increases hevin levels in human CSF samples. (A, B) ELISA analysis showing increased hevin levels in human CSF 10 hours after neurosurgery. (C) BCA protein assay showing total protein level increased in human CSF 10 hours after neurosurgery. (D) Fold changes of normalized hevin level in human CSF 10 hours after neurosurgery, normalized to total protein changes. n = 10 patients. \*\**P* < 0.01, \*\*\**P* < 0.001, paired t-test, Data shown as mean ± SEM.

## Discussion

Despite recent progress in demonstrating “gliopathy” in the pathogenesis of pain (15) and the previous studies showing an essential role of spinal TSP4 and cortical TSP1 in neuropathic pain sensitization (22, 23, 32), our knowledge is still limited regarding the specific mediators secreted by astrocytes that can drive neuropathic pain. We employed both loss-of-function and gain-of-function approaches to demonstrate a critical role of astrocytic hevin in driving neuropathic pain. Our data showed that nerve injury-induced mechanical allodynia in the maintenance phase, but not the induction phase, is impaired after hevin deficiency, in further support of the notion that astrocytes are crucial for the maintenance of neuropathic pain (15). Nerve injury induces marked and long-lasting astrogliosis in SDH that is correlated with the time course of neuropathic pain (6, 18). It has been shown that hevin is expressed by reactive astrocytes in the brain (33). We demonstrate that re-expression of hevin in SDH reactive astrocytes in hevin-KO mice is sufficient to reinstate neuropathic pain after nerve injury. Furthermore, intrathecal injection of hevin produced persistent mechanical allodynia in WT mice of both sexes.

Mechanistically, we demonstrated that hevin induces central sensitization and mechanical pain through regulation of GluN2B containing NMDARs in SDH neurons, in agreement with our previous report that hevin is crucial for the formation of thalamocortical connections in the visual cortex via specific regulation of GluN2B during development (8). Thus, hevin-induced mechanical pain was completely blocked by the GluN2B antagonist RO25. As expected, hevin deficiency resulted in a significant reduction of NMDA currents in SDH neurons through adulthood. Interestingly, physiological pain in adult mice at the baseline is unaltered in hevin-KO mice, suggesting a specific contribution of NMDARs to synaptic plasticity in pathological pain (3). Although we did not find significant change in AMPA-induced currents in SDH neurons of hevin-KO mice, a recent study shows that spinal hevin also mediates membrane trafficking of GluA1-containing AMPA receptors in remifentanil-induced postoperative hyperalgesia (34). Our findings strongly suggest that hevin-induced synaptic plasticity not only occurs during development but also manifests in pathological and neurodegenerative conditions such as neuropathic pain.

Our data also showed that hevin is highly secreted in mouse CSF and this secretion is further increased after nerve injury. Notably, CSF-HEVIN changes are clinically relevant. We found a rapid and substantial increase of HEVIN in human CSF samples 10 hours after painful neurosurgical procedures (31). Neurosurgery/otolaryngology procedures are also associated with an increase in the CSF tau levels that have been implicated in the development of dementia (31). It is of great interest to examine CSF-HEVIN levels in a general surgery population. Thus, CSF HEVIN could potentially serve as a biomarker for pain following neurosurgery or general surgery, an important question for evaluation in future studies.

It remains to be identified whether CSF-HEVIN is derived from astrocytes. It is well-established that overactivation of GluN2B causes neurotoxicity (35). Thus, high levels of HEVIN production and secretion in the CNS is not only associated with pain but may also underlie surgery-induced neurocognitive injury, such as delirium (36) by producing GluN2B-mediated neurotoxicity. Future studies are warranted to evaluate the involvement of hevin in neurocognitive function under physiological and pathological conditions. Importantly, we have demonstrated that neutralizing secreted hevin with a monoclonal antibody could effectively alleviate inflammatory and neuropathic pain. Thus, targeting secreted hevin may offer new therapeutics to manage chronic pain, post-operative neurocognitive injury, and neurogenerative diseases.

## Acknowledgements

This work was supported by Duke University Anesthesiology Research Fund and partially supported by NIH grants R01 grants DE17794 and DE29342 and DoD grant W81XWH2110885. MB acknowledges additional support from NIH grants K76-AG057022, P30-AG028716, and P30-AG072958. JPM was supported by NIH grants R01 HL130443 and AG074185. CE and JJR were supported by NIH grants R01 DA031833 and NS102237. CE is a Howard Hughes Medical Institute Investigator.

## Materials and Methods

### Reagents

We purchased formalin, carrageenan, NMDA from Sigma-Aldrich, TCN201 (GluN2A antagonist, Cat. No.4154), Ro25-6981 (GluN2B antagonist, Cat. No.1594) from Tocris, and recombinant hevin from RD system (Cat: 4547-SL). Purified recombinant hevin and hevin-ΔDE (lacking a.a. 351-440) proteins, anti-hevin monoclonal antibody 12:155, and anti-hevin monoclonal antibody 12:54 were from laboratory of Cagla Eroglu at Duke University Medical Center.

### Mice

Hevin KO mice were from Eroglu’s laboratory (9), and control SVE/129 mice (aged 8-12 weeks of both sexes) were obtained from Taconic Biosciences. These mouse lines were maintained at Duke University animal facility. We also used CD1 mice (Charlese River, 8-10 weeks) of both sexes for comparing hevin’s effects on mechanical pain. All animals were housed under a 12-hour light/dark cycle with food and water available ad libitum. Sample sizes were estimated based on our previous studies for similar types of behavioral and electrophysiological analyses (5, 19). Animals were randomly assigned to each experimental group. All the animal procedures were conducted in accordance with the National Institutes of Health Guide for the Care and Use of Laboratory Animals and approved by the Institutional Animal Care & Use Committee (IACUC) of Duke University.

### Mouse models of pain, CSF collection, and intrathecal injection

To produce inflammatory pain, diluted formalin (5%, 20 μl) or carrageenan (1.5 %, 20 μl) was injected into the plantar surface of a hindpaw (5). Neuropathic pain was induced by chronic constriction injury (CCI) as previously published (19). In brief, after the left sciatic nerve was exposed, three ligatures (LOOK® 6-0 Silk, #SP102) were placed around the nerve proximal to the trifurcation with one millimeter between each ligature. The ligatures were loosely tied until a short flick of the ipsilateral hind limb was observed. Mice in the sham group received surgery identical to those described but without nerve ligation. Mouse cerebrospinal fluid (CSF) was collected from the cisterna magna 14 days after CCI (30). For intrathecal injection, spinal cord puncture was made with a 30G needle between the L5 and L6 level to deliver reagents (10 μl) to the cerebral spinal fluid (30).

### Spinal injection of AAV2/5 virus

As our previous report (9), full-length hevin cDNA was cloned into the pZac2.1 gfaABC1D-Cyto-GCaMP3 viral vector (Addgene plasmid #44331), between the NheI and NotI restriction sites, replacing the GCaMP3 sequence. AAV2/5 particles were packaged and synthesized by the Penn Vector Core facility (University of Pennsylvania). Intra-spinal injections were performed as described previously (37). In brief, Hevin KO mice were deeply anaesthetized by subcutaneous (s.c.) injection of ketamine (100 μg/g) and xylazine (10 μg/g). Paraspinal muscles around the left side of the interspace between Th13 and L1 vertebrae were removed, and the dura mater and the arachnoid membrane were carefully incised using the tip of a 30G needle to make a small window to allow a glass micropipette (diameter of 80 μm) insert directly into the spinal cord dorsal horn (SDH). The microcapillary was inserted with AAV2/5 virus solution through the small window (approximately 500 μm lateral from the midline) and inserted into the SDH (250 μm in depth from the surface of the dorsal root entry zone). Each mouse was given single injection (0.7 μl, 1 x 10^12^ GC/ml) of AAV5.GFAP.Hevin-MycHis.SV40 virus or AAV5.GFAP.HevinDDE-MycHis.SV40 virus.

### Behavioral testing

Animals were habituated to the environment for at least 2 days before the testing. The room temperature and humidity remained stable for all experiments. All the behaviors were tested blindly. For testing spontaneous pain in formalin model, we measured the time (seconds) mice spent on licking or flinching the affected paw every 5 minutes for 45 minutes. For testing mechanical sensitivity, we confined mice in boxes placed on an elevated metal mesh floor and stimulated their hindpaws with a series of von Frey hairs with logarithmically increasing stiffness (0.02-2.56g, Stoelting), presented perpendicularly to the central plantar surface. We determined the 50% paw withdrawal threshold by Dixon’s up-down method (38). Thermal sensitivity was assessed by tail immersion, hot plate, and radiant heat tests. For tail immersion test, the lower 5.0 cm portion of the tail was dipped in hot water at 48 or 52 °C and the tail flick latency was recorded with a cut-off time of 20 seconds. For hot plate test, mice were placed on the hot plate at 50 or 56°C and the reaction time was scored when the animal began to exhibit signs of pain avoidance such as jumping or paw licking. Animals that did not respond to the noxious heat stimulus after 30 s were removed from the plate. For the radiant heat test, Hargreaves apparatus (IITC Life Science) was used and the basal paw withdrawal latency was adjusted to 9-12 s, with a cutoff of 20 s to prevent tissue damage.

Spontaneous and ongoing pain was assessed by conditioned place preference (CPP) test. We used a single trial conditioning protocol to measure CPP as previously report (27, 39). All mice underwent a 3-day pre-conditioning habituation and animal behavior was video-recorded. Analyses of the pre-conditioning (baseline) behavior showed no pre-existing chamber preference. On the conditioning day, mice received the vehicle (PBS, 10 μl, i.t.) control paired with a randomly chosen chamber in the morning, and clonidine (10 μg in 10 μl PBS, i.t.) paired with the other chamber 4 h later. Chamber pairings were counterbalanced. On the test day (3 w after CCI), 20 h following the afternoon pairing, mice were placed in the CPP box with access to both chambers and the behavior was recorded for 15 min and analyzed by ANY-maze software for chamber preference.

### Immunofluorescence and Imaging

After appropriate survival times after nerve injury, mice were deeply anesthetized with isoflurane and perfused through the ascending aorta with PBS and then followed by 4% paraformaldehyde. The L4–L5 spinal cord segments and L4-L5 DRGs were removed and postfixed in the same fixative overnight. Spinal cord (30 μm, free floating) and DRG tissue sections (14 μm) were cut in a cryostat. The sections were blocked with 2% goat or donkey serum for 1 h at room temperature and then incubated overnight at 4°C with the following primary antibodies: anti-hevin (rat anti-hevin monoclonal 12:155, 1 μg/mL, from Eroglu’s lab), anti-NeuN (mouse, 1:1000, Millipore, Catalog: MAB377), anti-GFAP (mouse, 1:1000, Millipore, catalog #MAB360), anti-IBA-1 (rabbit, 1:1000, Wako Chemicals, catalog #019-19741), and anti-Myc (mouse, 1:1000, Cell Signaling, Catalog: #2276). Tissue sections were then staining by cyanine 3(Cy3)-, cyanine 5(Cy5)- and/or FITC-conjugated secondary antibodies (1:400; Jackson ImmunoResearch) for 2 h at room temperature. The stained sections were examined with a Nikon fluorescence microscope, and images were captured with a CCD Spot camera. For high resolution images, sections were also examined under a Zeiss 510 inverted confocal microscope.

### Patch clamp recordings in spinal cord slices

As previously reported (19), we removed a portion of the lumbar spinal cord (L4-L5) from young mice (4-6 weeks old) under urethane anesthesia (1.5 −2.0 g/kg, i.p.) and kept the spinal cord segments in pre-oxygenated ice-cold artificial cerebrospinal fluid (ACSF). Transverse slices (400 μm) were made on a vibrating microslicer and the slices were transferred to ACSF saturated with 95% O_2_ and 5% CO_2_ at 26°C for at least 1 h prior to experiment. The ACSF contains (in mM): NaCl 125, KCl 2.5, CaCl_2_ 2, MgCl_2_ 1, NaH_2_PO_4_ 1.25, NaHCO_3_ 26, and glucose 25. The whole cell patch-clamp recordings were conducted from lamina IIo neurons in voltage clamp mode. Lamina IIo interneurons are predominately excitatory, make direct connections with lamina I projection neurons, and play critical role in transmitting mechanical pain (28, 29). Patch pipettes were prepared from thin-walled, borosilicate, glass-capillary tubing (1.5 mm o.d., World Precision Instruments) and filled with an internal solution containing following (in mM): K-gluconate 135, KCl 5, CaCl_2_ 0.5, MgCl_2_ 2, EGTA 5, HEPES 5, and ATP-Na_2_ 5. Data were acquired using an Axopatch 700B amplifier and were low pass-filtered at 2 kHz and digitized at 5 kHz. NMDA currents were recorded in IIo neurons by perfusing spinal cord slices with 100 μM NMDA for 3 sec in ACSF with TTX (5 μM). To measure NMDAR-mediated evoked EPSCs (NMDA-eEPSCs) in lamina IIo neurons, dorsal root (~ 5 mm) was stimulated through a suction electrode. NMDA-eEPSCs were pharmacologically isolated in Mg^2+^-free ACSF containing CNQX (10 μM), strychnine (2μm) and picrotoxin (100 μM). Neurons were voltage clamped at −40 mV and NMDA-eEPSCs were evoked at 0.05 Hz. QX-314 (5 mM) was added to the pipette solution to prevent discharge of action potentials. Synaptic strength was quantified by the peak amplitudes of eEPSCs. Data were collected with pClamp version 10 software and analyzed using Clampfit version 10.

### Western blot

SDH protein samples were prepared as we previously demonstrated (40), and 30 μg of proteins were loaded for each lane and separated on SDS-PAGE gel (4–15%; Bio-Rad). After the transfer, the blots were incubated overnight at 4°C with anti-hevin (rat anti-hevin monoclonal 12:155, 1 μg/mL). These blots were further incubated with HRP-conjugated secondary antibody, developed in ECL solution (Pierce), and the chemiluminescence was revealed by Bio-Rad ChemiDoc XRS for 1–5 min.

### Human CSF collection

Human CSF was collected from the patients enrolled in the Markers of Alzheimer’s disease after Propofol vs. Isoflurane Anesthesia (MAD-PIA) trial. MAD-PIA was a prospective randomized trial registered with http://www.clinicaltrials.gov (NCT01640275) on June 20, 2012, by Miles Berger, the study PI, with approval by the Duke University Institutional Review Board (31). Human CSF samples (10 ml) were obtained from the lumbar drain (AccuDrain INS-8400; Integra Neurosciences, Plainsboro, NJ, USA) at the time of drain placement (0 h) and 12 h after intracranial surgery.

### ELISA

Mouse hevin ELISA kit was purchased from LifeSpan BioSciences (Catalog: LS-F12654). Human hevin ELISA kit was from Cloud-Clone Corp (Catalog: SEM267Hu). ELISA was conducted in mouse CSF samples (10 μl) and human CSF (10 μl), according to manufacturer’s instructions. The standard curve was included in each experiment.

### Statistical analyses

All the data were expressed as mean ± s.e.m, as indicated in the figure legends. Statistical analyses were completed with Prism GraphPad 8.0. Biochemical and behavioral data were analyzed using two-tailed student’s t-test (two groups, unpaired or paired) or Two-Way ANOVA followed by post-hoc Bonferroni test. Electrophysiological data were tested using one-way ANOVA (for multiple comparisons) or Two-Way ANOVA (for multiple time points) followed by post-hoc Bonferroni test or student’s t-test (two groups, unpaired or paired). The criterion for statistical significance was *P* < 0.05.

---

**fig. S1.**
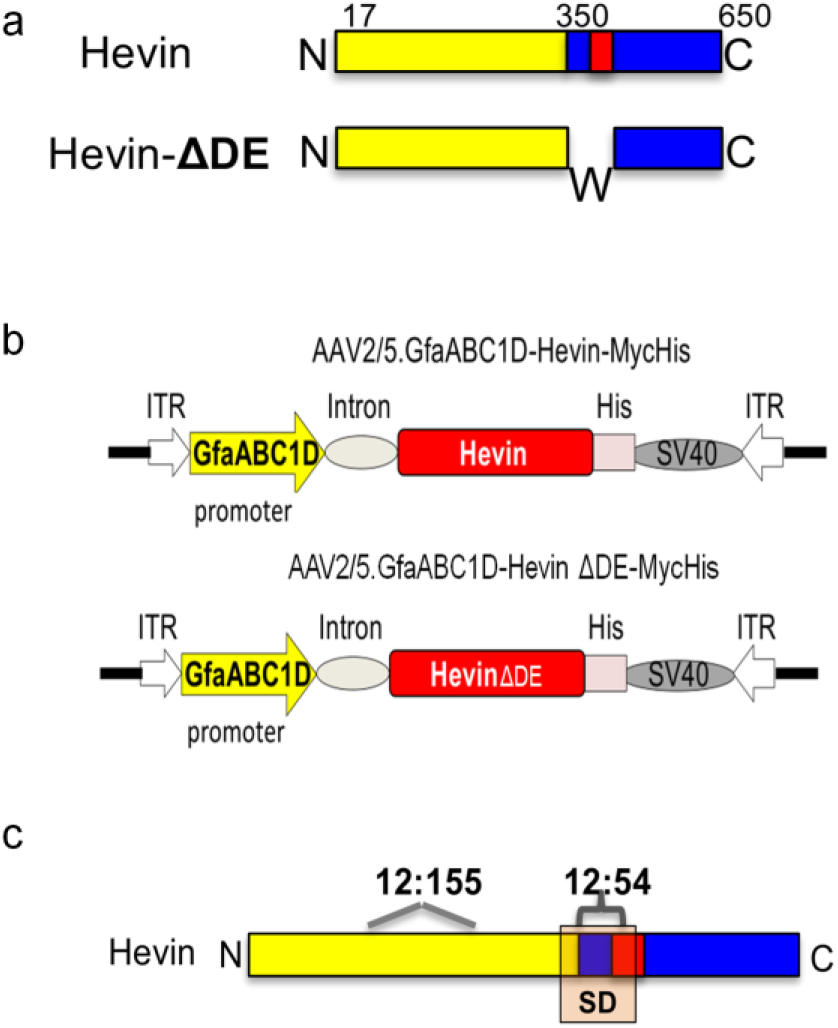
Schematic of hevin protein (a), hevin-AAV (b), and mapped epitopes (c) for anti-hevin monoclonal antibodies.

**fig. S2.**
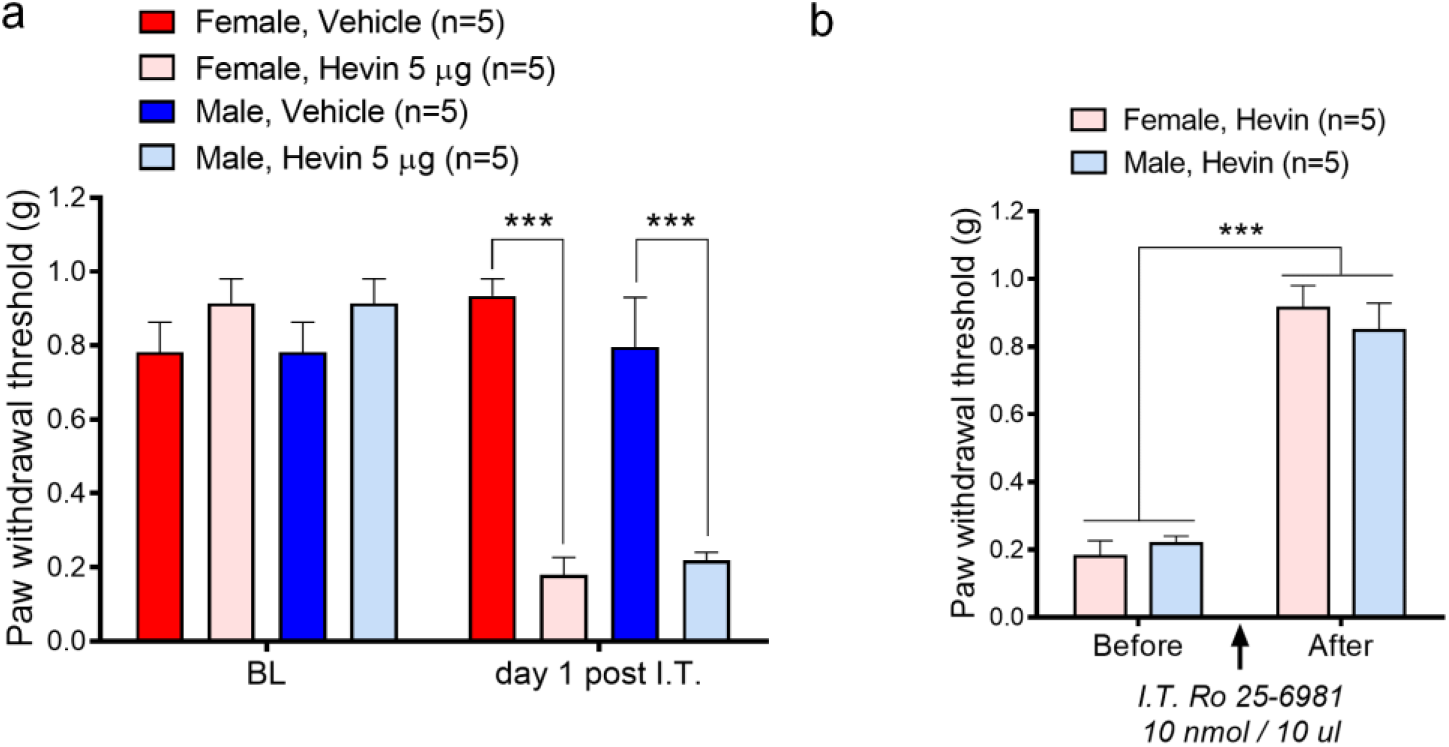
Hevin and NR2B activation drive mechanical pain in both sexes. (a) Intrathecal injection of purified WT hevin (5 μg, i.t.) induced robust mechanical allodynia in both WT naïve male and female mic for more than 1 day. n = 5 mice per group. \*\*\**P*<0.001, Two-Way ANOVA followed by Bonferroni posthoc test. Data shown as mean ± SEM. (b) Intrathecal hevin-induced mechanical allodynia was completely reversed by Ro25-6091 (GluN2B antagonist) in both WT male and female mice. n = 5 mice per group. \*\*\**P*<0.001. Two-Way ANOVA followed by Bonferroni posthoc test. Data shown as mean ± SEM.

**fig. S3.**
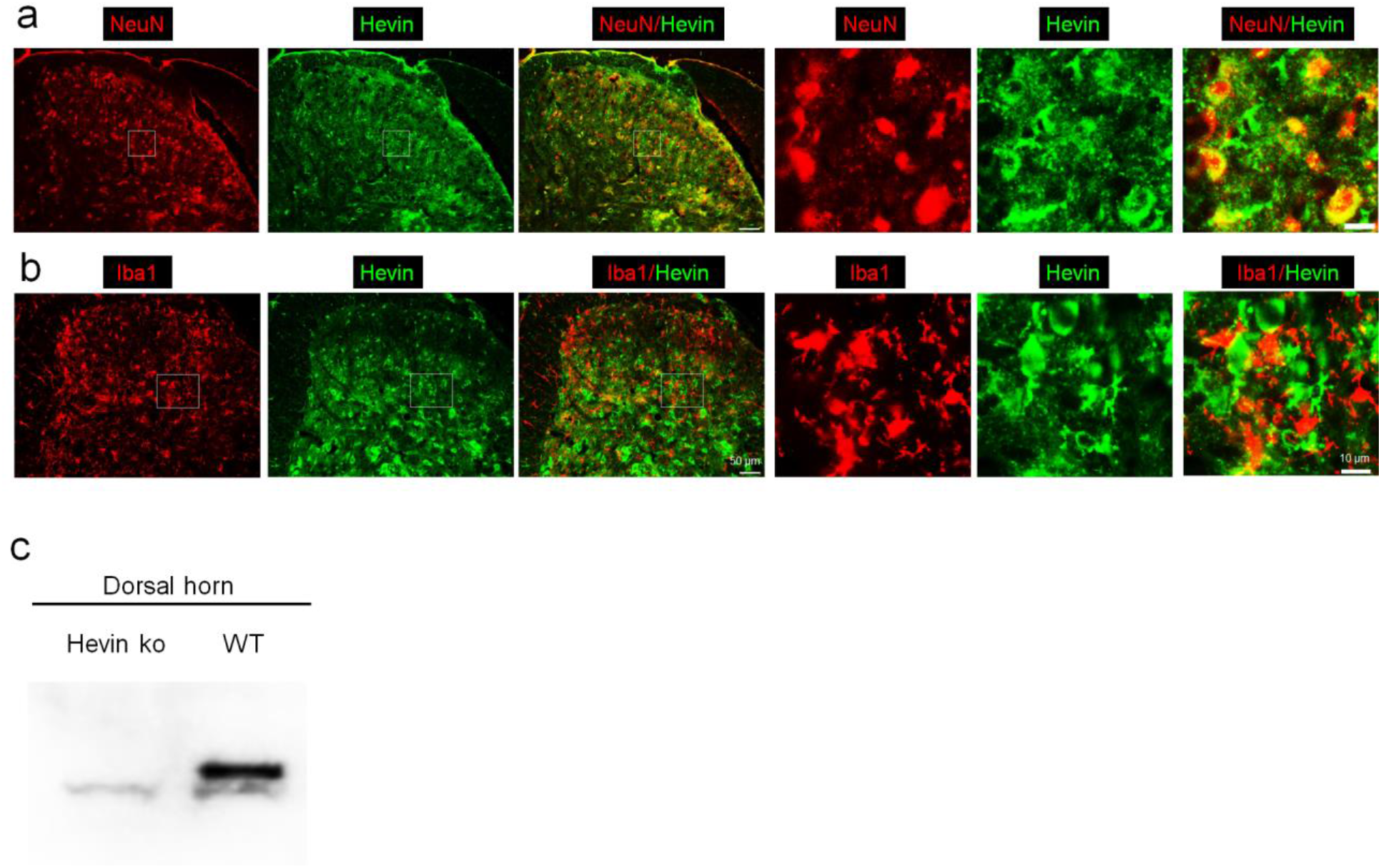
Hevin expression in SDH of WT and KO mice. (a, b) Double immunostaining of hevin (green) and the neuronal marker NeuN (red, a) or the microglial marker (Iba1, b) in the spinal cord dorsal horn in mice. Note hevin expressed by some neurons (NeuN^+^) but not microglia (Iba1^+^). Right images are enlarged images of the left image. Scale bars: 50 μm (left) and 10 μm (right). (c) Western blot results showed that hevin was expressed in spinal dorsal horn of WT mice and absent in KO mice.

**fig. S4.**
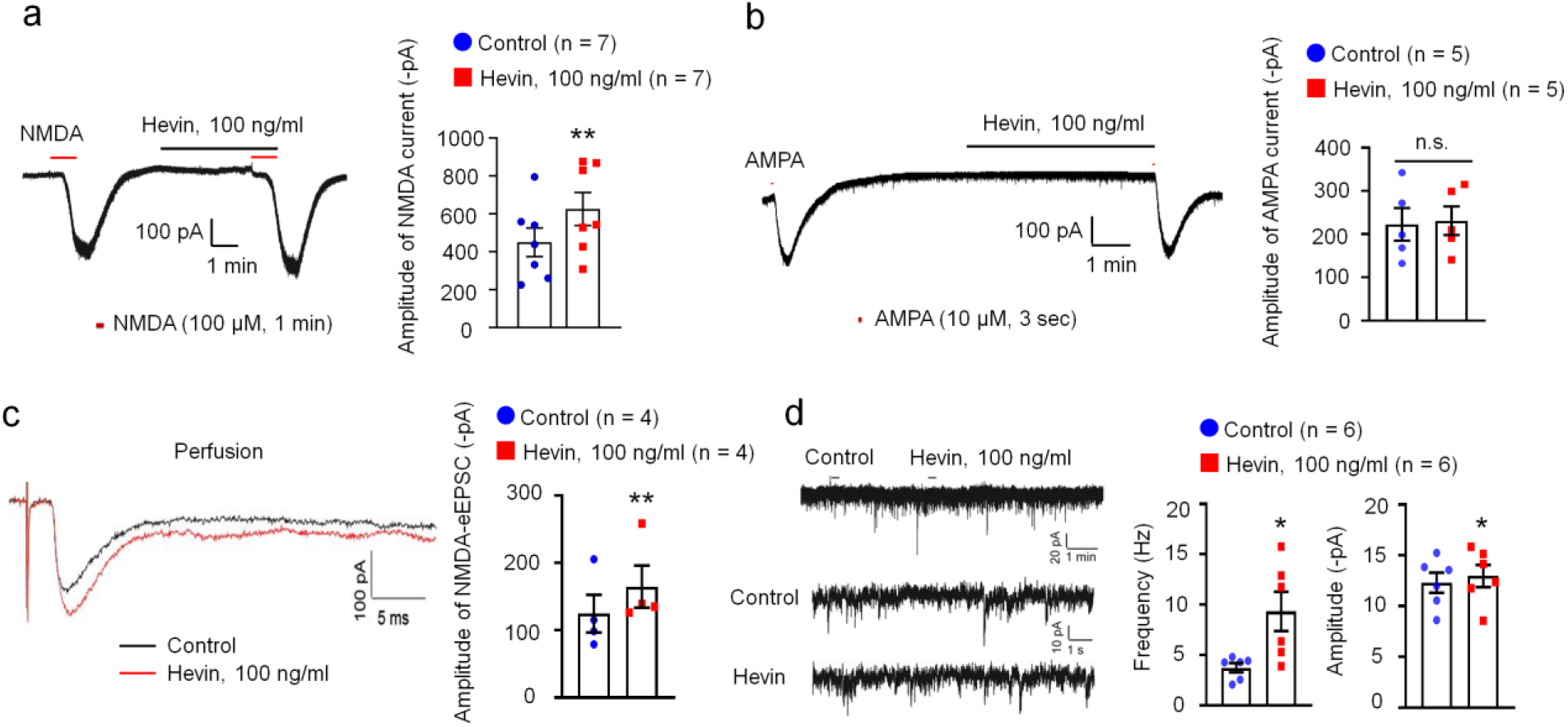
Electrophysiological characterization of hevin’s actions in lamina IIo neurons of spinal cord slices. (a, b) Acute effects of hevin (100 ng/mL, 5 min) on NMDA or AMPA-induced current in lamina II neurons. Note hevin enhanced the NMDA-induced currents but no effects on AMPA-evoked currents in lamina IIo neurons. n = 7 or 5 neurons per group (shown in each column). *\*\*P* < 0.01, paired t-test, n.s., not significant. Data shown as mean ± SEM. (c) Effect of hevin (100 ng/mL) on NMDA-eEPSC in lamina IIo neurons evoked by DR stimulation. Note hevin also enhanced the NMDA receptor–mediated eEPSCs. n = 4 neurons per group. \*\**P* < 0.01, paired t-test. Data shown as mean ± SEM. (d) Hevin superfusion (100 ng/ml) increased spontaneous EPSC frequency and amplitude in lamina IIo neurons of spinal cord slices from CCI mice. n = 6 neurons per group. \**P*<0.05, paired t-test. Data shown as mean ± SEM.

## References

1. M. Costigan, J. Scholz, C. J. Woolf, Neuropathic pain: a maladaptive response of the nervous system to damage. Annu.Rev.Neurosci. 32, 1–32 (2009).

2. R. Baron, A. Binder, G. Wasner, Neuropathic pain: diagnosis, pathophysiological mechanisms, and treatment. Lancet Neurol. 9, 807–819 (2010).

3. C. J. Woolf, M. W. Salter, Neuronal plasticity: increasing the gain in pain. Science 288, 1765–1769 (2000).

4. X. J. Liu et al., Treatment of inflammatory and neuropathic pain by uncoupling Src from the NMDA receptor complex. Nat.Med. 14, 1325–1332 (2008).

5. G. Chen et al., beta-arrestin-2 regulates NMDA receptor function in spinal lamina II neurons and duration of persistent pain. Nat Commun 7, 12531 (2016).

6. R. R. Ji, C. R. Donnelly, M. Nedergaard, Astrocytes in chronic pain and itch. Nat Rev Neurosci 10.1038/s41583-019-0218-1 (2019).

7. J. C. Foley, S. R. McIver, P. G. Haydon, Gliotransmission modulates baseline mechanical nociception. Mol Pain 7, 93 (2011).

8. Q. Xu et al., Astrocytes contribute to pain gating in the spinal cord. Sci Adv 7, eabi6287 (2021).

9. S. K. Singh et al., Astrocytes Assemble Thalamocortical Synapses by Bridging NRX1alpha and NL1 via Hevin. Cell 164, 183–196 (2016).

10. H. Kucukdereli et al., Control of excitatory CNS synaptogenesis by astrocyte-secreted proteins Hevin and SPARC. Proc Natl Acad Sci U S A 108, E440–449 (2011).

11. J. A. Stogsdill et al., Astrocytic neuroligins control astrocyte morphogenesis and synaptogenesis. Nature 551, 192–197 (2017).

12. A. Araque, V. Parpura, R. P. Sanzgiri, P. G. Haydon, Tripartite synapses: glia, the unacknowledged partner. Trends Neurosci 22, 208–215 (1999).

13. C. Eroglu et al., Gabapentin receptor alpha2delta-1 is a neuronal thrombospondin receptor responsible for excitatory CNS synaptogenesis. Cell 139, 380–392 (2009).

14. K. T. Baldwin et al., HepaCAM controls astrocyte self-organization and coupling. Neuron 109, 2427–2442 e2410 (2021).

15. R. R. Ji, T. Berta, M. Nedergaard, Glia and pain: Is chronic pain a gliopathy? Pain S0304-3959(13)00330-8[pii];10.1016/j.pain.2013.06.022 [doi] (2013).

16. S. A. Liddelow et al., Neurotoxic reactive astrocytes are induced by activated microglia. Nature 541, 481–487 (2017).

17. W. Guo et al., Glial-cytokine-neuronal interactions underlying the mechanisms of persistent pain. J.Neurosci. 27, 6006–6018 (2007).

18. M. Tsuda et al., JAK-STAT3 pathway regulates spinal astrocyte proliferation and neuropathic pain maintenance in rats. Brain awr025[pii];10.1093/brain/awr025 [doi] (2011).

19. G. Chen et al., Connexin-43 induces chemokine release from spinal cord astrocytes to maintain late-phase neuropathic pain in mice. Brain 137, 2193–2209 (2014).

20. H. Zhang, S. Y. Yoon, H. Zhang, P. M. Dougherty, Evidence that spinal astrocytes but not microglia contribute to the pathogenesis of Paclitaxel-induced painful neuropathy. J.Pain 13, 293–303 (2012).

21. Z. Y. Zhuang et al., A peptide c-Jun N-terminal kinase (JNK) inhibitor blocks mechanical allodynia after spinal nerve ligation: respective roles of JNK activation in primary sensory neurons and spinal astrocytes for neuropathic pain development and maintenance. J.Neurosci 26, 3551–3560 (2006).

22. D. S. Kim et al., Thrombospondin-4 Contributes to Spinal Sensitization and Neuropathic Pain States. J Neurosci 32, 8977–8987 (2012).

23. J. Park et al., Central Mechanisms Mediating Thrombospondin-4-induced Pain States. J Biol Chem 291, 13335–13348 (2016).

24. Y. Kohro et al., Spinal astrocytes in superficial laminae gate brainstem descending control of mechanosensory hypersensitivity. Nat Neurosci 23, 1376–1387 (2020).

25. J. P. Girard, T. A. Springer, High endothelial venules (HEVs): specialized endothelium for lymphocyte migration. Immunol Today 16, 449–457 (1995).

26. X. J. Liu et al., Treatment of inflammatory and neuropathic pain by uncoupling Src from the NMDA receptor complex. Nat Med 14, 1325–1332 (2008).

27. T. King et al., Unmasking the tonic-aversive state in neuropathic pain. Nat Neurosci 12, 1364–1366 (2009).

28. J. Braz, C. Solorzano, X. Wang, A. I. Basbaum, Transmitting Pain and Itch Messages: A Contemporary View of the Spinal Cord Circuits that Generate Gate Control. Neuron 82, 522–536 (2014).

29. B. Duan, L. Cheng, Q. Ma, Spinal Circuits Transmitting Mechanical Pain and Itch. Neurosci Bull 34, 186–193 (2018).

30. G. Chen, C. K. Park, R. G. Xie, R. R. Ji, Intrathecal bone marrow stromal cells inhibit neuropathic pain via TGF-beta secretion. J Clin.Invest 125, 3226–3240 (2015).

31. M. Berger et al., The Effect of Propofol Versus Isoflurane Anesthesia on Human Cerebrospinal Fluid Markers of Alzheimer’s Disease: Results of a Randomized Trial. J Alzheimers Dis 52, 1299–1310 (2016).

32. S. K. Kim et al., Cortical astrocytes rewire somatosensory cortical circuits for peripheral neuropathic pain. J Clin Invest 126, 1983–1997 (2016).

33. S. Lively, I. Moxon-Emre, L. C. Schlichter, SC1/hevin and reactive gliosis after transient ischemic stroke in young and aged rats. J Neuropathol Exp Neurol 70, 913–929 (2011).

34. Z. Wang et al., Spinal hevin mediates membrane trafficking of GluA1-containing AMPA receptors in remifentanil-induced postoperative hyperalgesia in mice. Neurosci Lett 722, 134855 (2020).

35. M. Aarts et al., Treatment of ischemic brain damage by perturbing NMDA receptor-PSD-95 protein interactions. Science 298, 846–850 (2002).

36. M. Berger et al., The INTUIT Study: Investigating Neuroinflammation Underlying Postoperative Cognitive Dysfunction. J Am Geriatr Soc 67, 794–798 (2019).

37. Y. Kohro et al., A new minimally-invasive method for microinjection into the mouse spinal dorsal horn. Sci Rep 5, 14306 (2015).

38. W. J. Dixon, Efficient analysis of experimental observations. Annu Rev Pharmacol Toxicol 20, 441–462 (1980).

39. G. Chen, C. K. Park, R. G. Xie, R. R. Ji, Intrathecal bone marrow stromal cells inhibit neuropathic pain via TGF-beta secretion. J Clin Invest 125, 3226–3240 (2015).

40. T. Berta et al., Extracellular caspase-6 drives murine inflammatory pain via microglial TNF-alpha secretion. J Clin.Invest 124, 1173–1186 (2014).

